# Early transatlantic movement of horses and donkeys at Jamestown

**DOI:** 10.1101/2024.06.11.598569

**Authors:** William Timothy Treal Taylor, Nicolas Delsol, Vicky M. Oelze, Peter Mitchell, Leah Stricker, Michael Lavin, Akin Ogundiran, Lauren Hosek, Christina Isabelle Barrón-Ortiz, Olumide Ojediran, Diana Quintero-Bisono, Dane Magoon, Matthew E. Hill, Ariane E. Thomas, Anna Waterman, David W. Peate, Lorelei Chauvey, Stéphanie Schiavinato, Laure Tonasso-Calvière, Luis Borges, Aitor Brito-Mayor, Jonathan Santana, George Kamenov, Ludovic Orlando, John Krigbaum

## Abstract

Domestic horses and donkeys played a key role in the initial colonization of the Atlantic seaboard of the Americas, a process partially chronicled by historical records. While Spanish colonists brought horses to the Caribbean and southern latitudes earlier, the transport of domestic horses to the English colony at Jamestown, Virginia in 1606 was among the first dispersals to the eastern seaboard. Archaeozoological analysis, isotope analysis, and radiocarbon dating of identifiable domestic equid remains from two contexts associated with the initial occupation of Jamestown demonstrate intense processing and consumption of the first Jamestown horses during the “Starving Time” winter of 1609, while paleopathological data show evidence of their use in transport. Osteological, genetic, and isotopic study of these equid remains reveal the presence of at least one adult domestic donkey with mixed European and West African ancestry, possibly supplied through undocumented exchange during a trans-Atlantic stopover. These results reveal the importance of equids in the survival of early European settlers and the global connectivity of early trans-Atlantic exchange in horses and donkeys, showing Caribbean and African links in the founding livestock populations and pointing towards an important and ecologically-anchored role for donkeys in the early colonial lifeways along the Eastern seaboard.

## Introduction

The introduction of domestic horses into North America was among the most consequential events in the history of the Americas, with horses both transforming the lifeways of many Indigenous people as well as forming a core part of colonial infrastructure for European colonists [1]. Although the first horses reached areas of western and southeastern North America through the activity of Spanish explorers and settlers, many of the first domestic equids to reach the eastern seaboard of the United States were those that accompanied English settlers. During later centuries, the expansion of these British colonial horses would come to shape the genetic landscape of both colonial and Indigenous horse populations across the continent [2]. Jamestown, the first permanent English settlement in North America, represents an important snapshot in the early chapters of the human-equid story of the Americas.

The colony of Jamestown, located near the confluence of the James River with Chesapeake Bay in the present-day state of Virginia, was initially settled by 104 men and boys who sailed in a fleet of three ships from London in 1606. First-person accounts document this voyage, which stopped in the Canary Islands, crossed the Atlantic to the Caribbean and made numerous stopovers throughout (Martinique, Dominica, Nevis, St. Croix, Puerto Rico, and the Mona and Monito Islands) before arriving at their final destination [3,4]. A first and second supply voyage arrived in January and September 1608, respectively, likely following a similar route. In June 1609, the Third Supply voyage departed England for Jamestown, arriving in August, but a hurricane stranded one of seven ships from the fleet in Bermuda, where it spent the winter. Documentary evidence for the first dispersal of English horses to the colony comes from Gabriel Archer, who, when arriving at Jamestown on the *Blessing* in late August 1609, wrote that “sixe Mares and two horses” were taken aboard in Plymouth, England, before setting sail [5]. John Smith recorded that upon his departure from the colony in October 1609 Jamestown had by his count, “six mares and a horse,”suggesting that one of the horses perished during the journey [3].

Initially managing tenuously peaceful relations with Tsenacomocoan people, relations soured by the winter of 1609, resulting in a siege of the palisaded James Fort. This winter, known as the “Starving Time”, introduced extraordinary subsistence difficulties, and most of the colonists starved to death or perished from disease. Contemporaneous writings chronicle decisions to slaughter both livestock and animals not otherwise eaten, including horses, before eating dogs, rats, and even their deceased compatriots - although dogs were also consumed before the Starving Time [6]. In the spring of 1610, a beleaguered resupply expedition reached the settlement, discovering only 60 survivors. The lieutenant governor, Sir Thomas Gates, attempted to resuscitate the colony by establishing martial law, but only weeks later decided to abandon the fort. However, arriving on the Fourth Supply from England, Governor Lord De La Warr met Gates and his ships only about 10 miles into their departure, forcing them to turn back. Upon their return, De La Warr reinforced Martial Law and instituted a cleansing and rebuilding effort of James Fort. Eventually, supply ships would arrive to re-establish livestock populations, including domestic horses. However, beyond their initial transport to the settlement and their demise, little is known about the role of domestic horses and potentially other equids, such as donkeys and mules, as Jamestown first developed.

Archaeological animal remains from two features linked to the Starving Time provide a unique opportunity to understand the role of domestic equids in early British settlement of the eastern seaboard of North America. The first, excavated by Jamestown Rediscovery archaeologists in 2009, was a large rectangular cellar (‘Kitchen Cellar’) and barrel-lined well (‘First Well’) located in the geographic center of the fort. Filled with over half a million artifacts, including faunal remains, the feature is likely the colony’s first well, constructed in late 1608-early 1609 on the orders of President of the colony, John Smith [3]. Designated JR2718/Structure 185, artifacts and stratigraphic relationships within the well feature suggest that the well was filled during the De La Warr cleansing in the spring of 1610, with successive filling episodes occurring in later years of the 17th century as the fill settled [7]. Preliminary faunal analysis of material recovered from the well feature suggests a reliance on local species, including, fish, crab, turtle, wild bird, and wild small mammals in addition to species likely consumed during the Starving Time winter, including horse and dog remains [6]. Just to the east of the First Well is a second, likely contemporaneous feature, excavated by Jamestown Rediscovery archaeologists in 2012. Designated JR3081/Structure 191, the feature was an L-shaped subterranean cellar with two brick-facade ovens. Likely the colony’s first kitchen described in early records as constructed in early 1608, archaeological evidence suggests this feature was abandoned during the Starving Time winter and like the First Well, was filled with trash in the spring of 1610 [7]. Faunal analysis of material recovered from the Kitchen Cellar supports this interpretation [8]. Prior faunal analysis of the assemblage from these two locations revealed a total of 77 specimens provisionally identified as “horse.”

To assess the role of domestic equid use during the earliest occupation at Jamestown, we conducted interdisciplinary archaeological, osteological, and biomolecular (radiocarbon dating, isotopes, and DNA) analysis of equid bones linked to Starving Time contexts from Jamestown.

## Results

### Radiocarbon dating

Direct radiocarbon dating of five equid specimens produced three successful radiocarbon dates, including two from adult domestic horses (117294, 121460), and one from a specimen identified as an adult domestic donkey (121161). After calibration, each of these specimens produced a nearly identical radiocarbon measurement consistent with an animal that died during the 16^th^ or early 17^th^ century (Table 1). Combining these three measurements in a simple Bayesian uniform phase model, and assuming that the deposition of the horses must have postdated the first known arrival of European horses on the North American mainland with Ponce de León, the modeled onset of equid use at these two features is dated to between 1577-1610 CE (1 sigma calibrated range), with a median date of 1592 CE. The termination of equid activity at these two features is modeled at ca. 1601-1634 cal. CE (1 sigma), with a median modeled date of 1619 CE. These results are therefore consistent with the archaeological context and the assumption that the two features date to the Starving Time winter of 1609-1610, particularly when we consider that the horse and donkey tooth dentin, that was sampled for dating, likely formed during tooth mineralization several years before the animals’ death.

### Elements present

Skeletal elements present in the assemblage include specimens from both appendicular and axial portions of the skeleton, including teeth, vertebrae, limbs, and pelvis. Given the highly processed nature of the identified elements, and the lower caloric value of many of them (such as the bones of the lower limbs), it is highly likely that elements of the entire skeleton were originally represented among the full assemblage, but that larger limb bones, ribs, and vertebrae have been processed so intensively as to render them difficult to identify as seen in other comparable assemblages [9]. Alternatively, it may be that these lower-value elements were disposed of in the two depositional contexts analyzed here, the cellar and the well, while high-value components were curated and/or discarded in a different manner.

### Species identification

For each element, species previously identified as *Equus* sp. were assessed against digitized comparative specimens from the Archaeozoology Laboratory at the University of Colorado, and we excluded those that could not be positively confirmed as *Equus* from further analysis. Of these confirmed as *Equus*, 62 specimens were consistent with an identification as domestic horse (*Equus caballus*) or mule (*Equus caballus x Equus asinus*), while one small upper cheek tooth (upper M2) was identified as domestic donkey (*Equus asinus*).

The occlusal surface of the tooth (catalog # 121161) identified as domestic donkey was sufficiently well preserved for further assessment (Figure 2A). This tooth shows dental features that are typically found in *Equus asinus*. In particular, the distal length of the protocone is approximately equal to the mesial length of the protocone (i.e., the protocone is symmetrical), the *pli caballin* is absent, and the post-protoconal groove is deep. These features are found at a higher frequency in domestic donkeys than in domestic horses (Supplementary Material Section 2). The five samples analyzed for their DNA were identified as four horse mares and one donkey jack, which confirmed the taxonomic status assigned morphologically (Supplementary Appendix 1).

**Figure 1.**
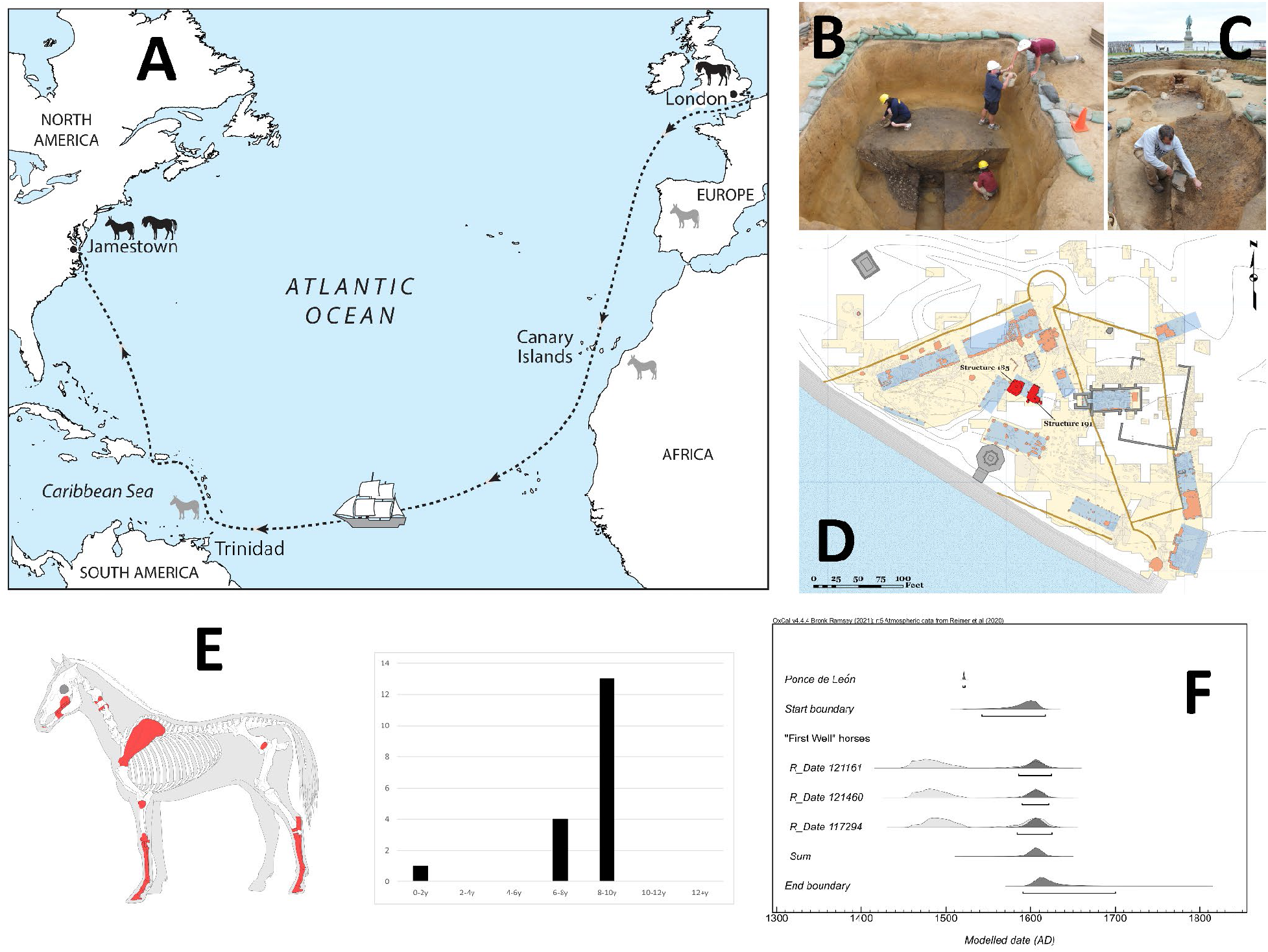
**A**, top left. Study sites and localities mentioned in the text, along with the route of the first colonial voyage to Jamestown. Drawing by Bill Nelson. **B**, top center. Excavation of the First Well at Jamestown, JR2718/Structure 185. **C**, top right. Excavation of the Kitchen Cellar, JR3081/Structure 191. **D**, center right. Site map with features highlighted. Analyzed structures are highlighted in red, with excavated areas in yellow and identified structures in blue, other archaeological features in orange. Location of the original 1607-08 fort palisades are outlined in goldenrod. **E**, bottom left. Skeletal elements identified and present in the analyzed assemblage (left), and estimated age of horse specimens in the assemblage that could be aged using dentition (right). **F**, bottom right. Bayesian uniform phase model for horse activity in Structures 185 and 191 associated with the Starving Time at Jamestown.

**Figure 2.**
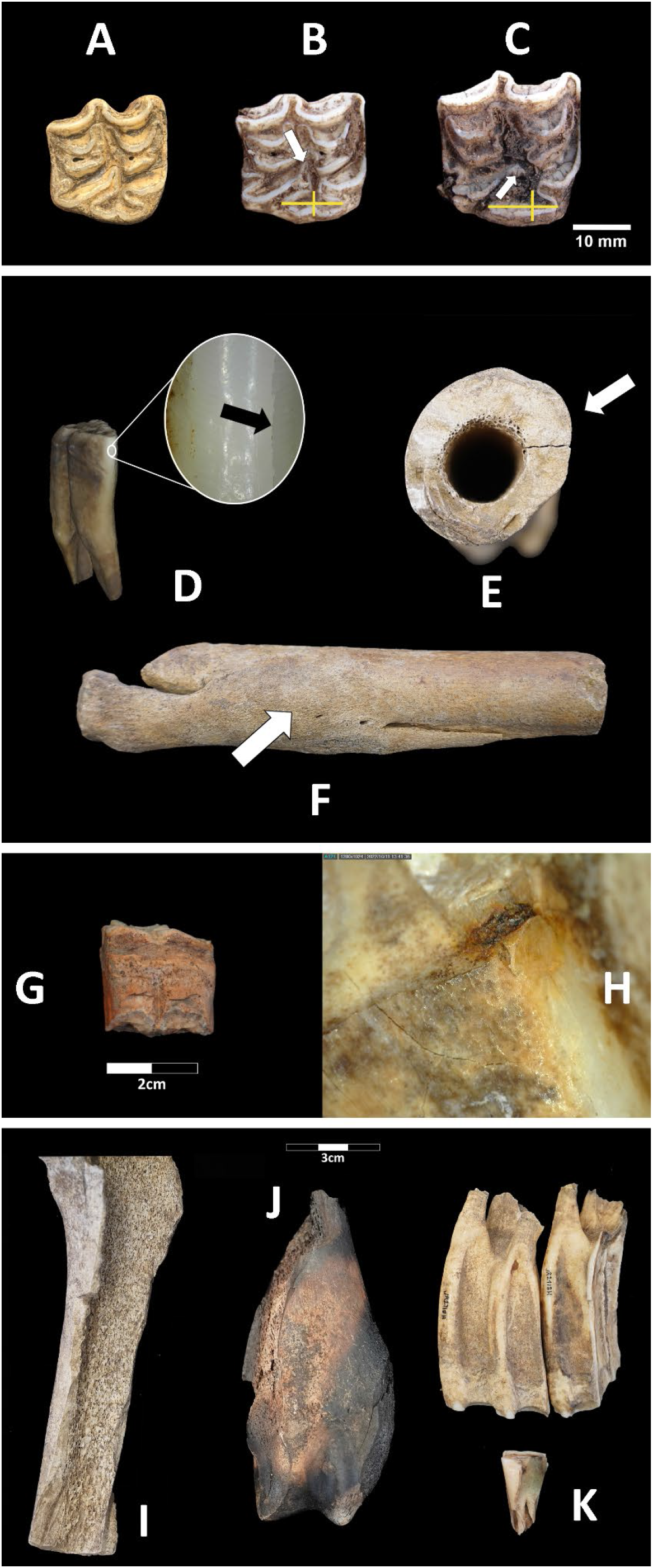
Occlusal view of Jamestown Equus asinus tooth (upper M2) and upper M2 molars of E. asinus and E. caballus. (**A**) Jamestown upper M2 identified as E. asinus. (**B**) upper M2 of E. asinus with symmetrical protocone (i.e., the distal length of the protocone is approximately equal to the mesial length of the protocone; yellow cross), pli caballin absent, and deep post-protoconal groove (white arrow). (**C**) upper M2 of E. caballus with asymmetrical protocone (yellow cross), pli caballin present (white arrow), and shallow post-protoconal groove. All teeth are upper right molars; mesial side to the right. **D**) Longitudinal fracture of the anterior enamel surface on a Jamestown horse, possibly linked with metal bit use. **E**) Cross-sectional asymmetry of a fractured metapodial element from Jamestown, suggestive of repeated activity patterns linked with transport. **F**) Healed splint bone fracture of a horse specimen from Jamestown. Transverse splitting of mandibular cheekteeth in the Jamestown assemblage (**G**), and embedded iron fragments indicative of axe use for splitting (**H** ). Modified Starving Time-era horse bones from Jamestown, including limb bones with removed trabecular bone (**I**) and burning (**J**), and teeth with evidence for longitudinal splitting and pulp extraction (**K**).

### Demographics

Age estimation of the identified horse specimens shows a profile of mostly so-called “prime age” adults between the ages of 6-10 years, with a Minimum Number of Individuals (MNI) of at least two adult animals. This finding, along with the DNA-based determination of all analyzed horse teeth as female, is consistent with the hypothesis that the analyzed assemblage was derived from the horses known from first-person accounts, consisting of 8 total animals, six females and two males. The estimated age of the specimens, with a central tendency of ∼8 years, would indicate that the animals were young adults near maturity (∼4 years) when they left England. We identified no canine teeth or other direct evidence of sex of the analyzed specimens but did identify the remains of a single juvenile specimen, a fractured phalanx of a young horse less than 1 year. The presence of this juvenile horse demonstrates that the first horses brought to the Jamestown settlement successfully reproduced at least once prior to the winter of 1609. Although age determination using techniques like crown height is not reliable on the basis of available data for donkeys [10], the donkey remains also provide an MNI of at least one adult donkey of greater than ∼2 years of age, based on its status as erupted and in wear with fully-formed roots [11].

### Pathology

Pathological features on the horse remains recovered from Starving Time features at Jamestown suggest that these animals were used in transport during their lifetime. Two second premolar specimens from the assemblage show exposure of the anterior enamel, a damage pattern often found in horses that have been ridden with a bit [12]. Although anterior damage of the lower second premolars is not uniquely indicative of bridle use [13], in one case the enamel is also fractured vertically in a manner consistent with traumatic contact from the anterior direction (Figure 2D), consistent with the use of a bridle mouthpiece. The exposed cross-sectional profile of one horse metatarsal shows asymmetry that could be linked to transport or work activities (Figure 2E), but may also reflect other aspects of activity patterns, such as gait [14,15].

Several fractures in the assemblage are likely associated with kick injuries. In one case, a left metacarpal bone shows a healed fracture of the splint bone (Figure 2F), which is commonly caused by kick injury from other animals. Similarly, the phalanx of the single foal identified in the assemblages exhibits a healed comminuted or crush fracture that reflects a high-energy injury [16]. These patterns could reflect animals being held together in close confinement in trans-Atlantic transport or in James Fort itself. One fragment of a mandible also showed an oval lytic lesion on the inferior lateral surface at the gonial angle; this feature likely represents a cyst with reactive bone formation along the floor of the cavity.

### Taphonomy

Horse and donkey remains from Jamestown exhibit an extraordinary level of cultural modification (Figure 2 G-K), with all but 11 (82%) of the analyzed specimens exhibiting some clear form of modification (spiral fractures (n = 6), cut marks (n = 13), chop marks (n = 20), burning (n =7), mid-shaft discoidal fractures (n =2), and cortical bone removal or pulp extraction (n = 22). Most striking is the degree to which all marrow-bearing elements have been processed for marrow and grease extraction, even those typically considered to be low-yield [17] up to and including the nearly-ubiquitous splitting of teeth to expose the pulp cavity (Figure 2G,H). Under low-power microscopy, some tooth fragments even displayed embedded pieces of iron apparently linked to ax use (Figure 2H). Finally, one horse tooth exhibited severe fluvial-like rounding that may represent “pot polish” or a mix of sheen, smoothing, and beveling/rounding that can be acquired during the process of boiling [18].

### DNA analysis

DNA preservation levels were compatible with the genome-wide characterization of the donkey jack specimen JT02/121161 (Jamestown, hereafter) (Supplementary Appendix 1). Neighbor-joining phylogenetic reconstructions supported the placement of the Jamestown specimen within a 100% bootstrap supported monophyletic group including all the previously-sequenced modern and ancient donkey specimens from Europe, the Canary Islands, and Brazil present in our comparative panel (Figure 3). Maximum Likelihood phylogenetic reconstructions in TreeMix confirmed general genetic affinities of European origins in both the Chupaderos (17^th^-19^th^ century Mexico[2]) and Jamestown specimens, regardless of whether groups of modern accessions were stratified by country of origins or subcontinental region (Figure 3C). This finding was consistent with the results of a principal component analysis, in which ancient European specimens, including those from Chupaderos and Jamestown, projected onto present-day European variation, and towards positive values along the first component, which is characteristic of modern European populations (Figure S3,S4). In this analysis, the placement of the Chupaderos donkey was closer to the modern populations from mainland Spain (ESP), the Canary Islands (CYK), and Brazil (BRA), relative to that of the Jamestown specimen. The latter was closer to the ancient European donkey from western Europe, including medieval (Albufeira, Portugal: ca. 1228-1280 cal. CE) and post-medieval (Fiumarella, Italy: ca. 1683-1936 CE), as well as from Antiquity (Boinville, France: ca. 200-500 CE, and; Bourse, France: ca. 0-500 CE).

**Figure 3.**
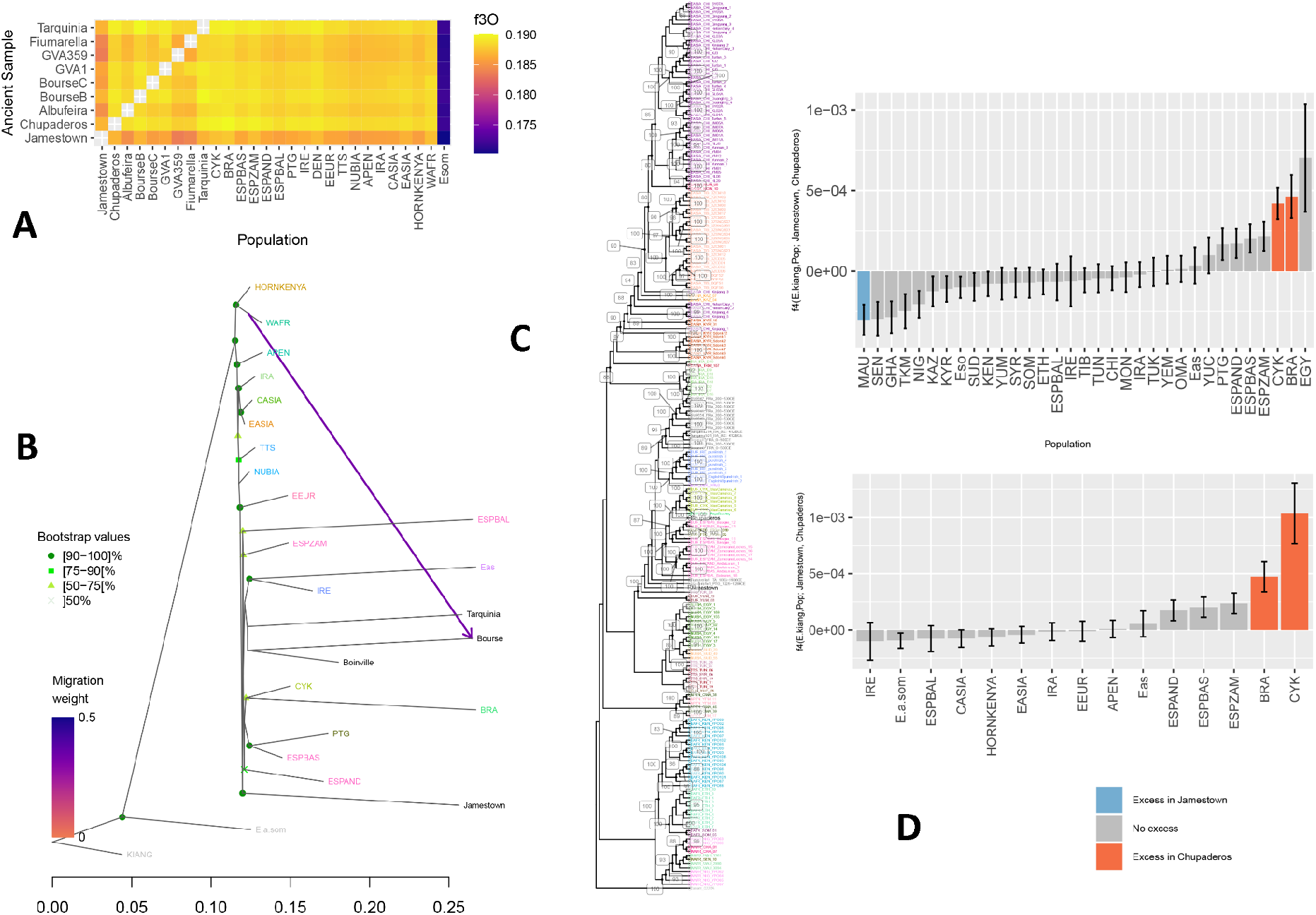
**A**, top left. f3-Outgroup statistics between a subset of ancient and modern donkeys. The group of Equus kiang genome sequences present in the genome panel served as the outgroup. The f3-outgroup statistics provides a measure for pairwise population genetic affinities. Modern donkey accessions were stratified according to the subcontinental groups of genetic affinities defined by Todd et al. [19], with Spanish accession subdivided to gain further resolution into possible genetic affinities in the region (ESPAND: Andalusia; ESPBAS: Basque; ESPZAM: Zamorano Leones, and; ESPBAL: Baleares). **B**, bottom left. Phylogenetic relationships between the ancient donkey specimen from Jamestown (JT02/121161) and a worldwide panel of modern donkeys, grouped by subcontinental groups of genetic affinities. The tree was rooted on the sequence data available for the Equus kiang species, and node supports were assessed from 100 bootstrap pseudo-replicates. The tree models population genetic affinities, considering an optimal number of migration edges (Supplementary Appendix 1). Subcontinental groups of genetic affinities are defined according to Todd et al. [19], with Spanish accession subdivided to gain further resolution into possible genetic affinities in the region (ESPAND: Andalusia; ESPBAS: Basque; ESPZAM: Zamorano Leones, and; ESPBAL: Baleares). **C**, center. Phylogenetic affinities between 223 donkey specimens distributed across the world. Node supports were estimated from 100 bootstrap pseudo-replicates and are provided in percentage. The labels of modern donkey accessions report the cluster of genetic affinities defined by Todd and colleagues [19], followed by the sample name. The labels of ancient donkey genomes indicate the sample name, followed by their country of origins (three letters code) and their estimated radiocarbon age and/or the timeline of their associated archaeological context, except for the Jamestown JT02/121161 and Chupaderos specimen from the 18^th^-20^th^ century CE. **D**, right 4-statistics of the form f4(Equus kiang, Pop; Jamestown, Chupaderos). This statistics measures whether the ancient donkey specimens from Jamestown and Chupaderos are genetically equally related to modern donkey accessions, grouped by country of origins (panel a), or stratified by the subcontinental groups of genetic affinities defined by Todd et al. [19] (panel b). Statistically significant positive (negative) values indicate extra-genetic affinities between Population (Pop) and Chupaderos (Jamestown).

Interestingly, the historical specimen from Chupaderos, Mexico, placed within a monophyletic group of modern donkeys from the Canary Islands, while the specimen from Jamestown did not group with any modern or ancient donkey specimen. This was consistent in both Neighbor-Joining and TreeMix phylogenetic reconstructions, and with f3-outgroup statistics, which measure pairwise genetic affinities between those samples and a comparative panel of other ancient donkeys from Europe, and worldwide modern donkey groups stratified by country, or subcontinent. The latter analyses revealed the different genomic makeup of both specimens, with Jamestown appearing closer to modern donkeys from the Balkan (YUM, North Macedonia) and Chupaderos also showing strong genetic affinities with Brazilian Pega donkeys and modern CYK donkeys. The presence of extra-genetic affinities with Pega donkeys and the Canary Islands in the Chupaderos specimen, relative to Jamestown, was confirmed by statistically significant f4-statistics of the form f4(*E. kiang*, X; Jamestown, Chupaderos), where X represented any modern donkey accession, grouped by country of subcontinental groups Figure S5). This analysis also revealed an excess of genetic sharedness between modern accessions from Western Africa (Mauritania), relative to Chupaderos. The presence of this extra-genetic affinity was no longer supported when grouping all modern accessions from Western Africa together (Mauritania as well as Senegal and Nigeria), suggesting specific contributions from Mauritania or closely related populations. This finding echoes the work from Todd and colleagues [19], which identified genetic contributions between Western Europe and Western Africa in the Roman and Medieval time periods, which were replicated in our own TreeMix phylogenetic reconstructions.

Finally, the genome sequenced from the Jamestown donkey was only compatible with one-way admixture modeling considering the Medieval specimen from Albufeira, Portugal, or the Chupaderos, Mexico, specimen as possible sources (Figure S5). Multiple two-way qpAdm models confirmed that the Jamestown genome consisted mainly of Portuguese (PTG, Portugal), Mediterranean (TUN, Tunisia, and; YUMYUC, North Macedonia and Croatia), the Levant (SYR, Syria), or Anatolia (TUK, Turkey) genetic ancestry, and approximately 8.4-13.4 % from Western African (MAU), in line with TreeMix analyses and f4-statistics (Figure 3). No such models included genetic contributions from mainland Spain or northern Europe (IRE, Ireland, and; DEN, Denmark). qpAdm modeling indicated that the general affinity of the Chupaderos genome and modern populations from Europe and the TUK population from Anatolia, as most one-way models could not be rejected. This was also true when considering the Roman specimens from Boinville, France, and the post-Medieval specimen from Fiumarella, Italy, as a possible source. Other ancient specimens from Europe included a larger range of possible population sources. This indicates donkey genetic profiles changing through space and time during the last 2,000 years in Europe, including variable degrees of Western African ancestry, with some ancient specimens having a more pervasive and widespread genetic legacy in the region today.

Combined, our analyses indicate that the donkey specimen excavated at Jamestown exhibited a genetic profile most characteristic of European ancestry, but also included Western African ancestry, as the best model in our dataset depicts Mauritania as a source. The latter genetic contribution was absent in the ancient specimen from Chupaderos, Mexico, which indicates different population sources for the origin of the two donkey specimens. While the specimen from Chupaderos appeared genetically close to the Pega donkey from Brazil, and modern donkeys from the Canary Islands, these specific genetic affinities were not found in the genome of the specimen from Jamestown.

### Isotopic Evidence on Diet and Possible Origins

The two teeth subjected to isotope analysis, a horse upper third molar (M3) and a donkey upper second molar (M2), represent sequences that mineralized at different developmental stages, with the horse M3 mineralizing during the third and fourth year (25 to 55 months [20]), and the donkey M2 beginning mineralization towards the end of the first year (8 to 18 months [11]).

The enamel δ^13^C values of the horse average -13.2‰ indicates a C_3_-plant based diet typical for temperate climate and vegetation zones whereas the donkey average enamel δ^13^C values of -8.8‰ suggests a mix of C_3_ and C_4_-plants, which could include wild C_4_-grasses but also domesticated C_4_-plants such as millet, sorghum and maize, found in warmer, dryer regions. This finding of broad differences in fodder is supported by the disparate δ^18^O values in the two equids. The horse exhibits an average δ^18^O value of -6.0‰, which is ^18^O-depleted compared to that of the donkey, with an average δ^18^O value of -2.5‰. This again strongly suggests different points of origin for these two equids.

However, the radiogenic isotope ratios of strontium and lead are similar among the two equids sampled, and the respective averaged ^87^Sr/^86^Sr ratios of 0.7104 and 0.7100 are found in many regions of the world. The averaged Pb ratios of the horse and donkey enamel samples show a narrow range, with ^206^Pb/^204^Pb ranging from 18.410 to 18.504, ^207^Pb/^204^Pb from 15.625 to 15.636, and ^208^Pb/^204^Pb from 38.367 to 38.449 (Table S2). These values are not typically found in the Americas [21] and likely derive from metal artifacts the two equids were exposed to either during life or postmortem diagenesis (Supplementary Material Section 4). Matching these isotope values to available isoscape resources for Britain [22] suggests that, while the ^87^Sr/^86^Sr and ^206^Pb/^204^Pb ratios of the horse are present in the biosphere of modern Britain (Fig. S1-A), its carbonate δ^18^O values converted to precipitation/drinking water δ^18^O values (-10.5‰) lie outside of the range of precipitation within Britain (Figure 4). Similarly, the average ^87^Sr/^86^Sr and ^206^Pb/^204^Pb ratios of the donkey match larger regions of Britain (Fig. S1-C), but only overlap with the donkey’s drinking water δ^18^O values in the far southwestern coast, and hence the warmest region of the UK (Fig. S1-D). This implies that both equids likely originated from warmer climatic regions. As an alternative, we matched the equine ^87^Sr/^86^Sr ratios and δ^18^O values to a novel bioavailable Sr isoscape available for larger parts of West Africa [23] as well as global precipitation δ^18^O data [24]. Results point to high probabilities of geographic origin for both equines in the coastal areas of Senegambia and present-day Sierra Leone (see Fig. S2).

**Figure 4.**
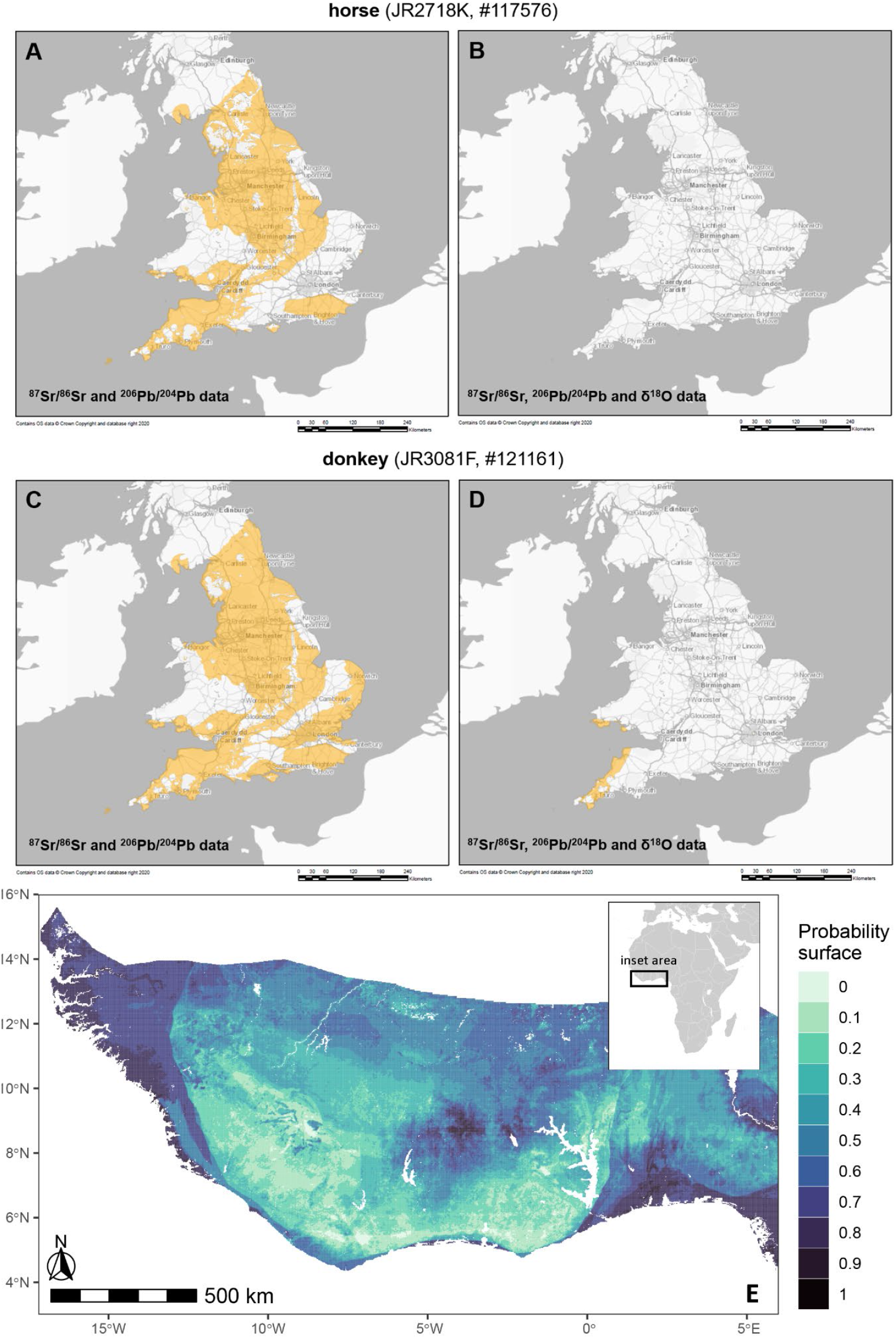
**A-D**. Matching the stable and radiogenic isotope data from a Jamestown horse and donkey to modern isotopic biosphere data from Britain [22] suggests that both animals match Britain in their ^87^Sr/^86^Sr and ^206^Pb/^204^Pb ratios, whereas the δ^18^O values do not. While the donkey’s δ^18^O values only match those of precipitation in the far southwest of the British Isles and suggest provenience in a warmer climate overall, those of the horse do not match modern precipitation in Britain at all, but suggest a much colder climate than modern day Britain, such as that prevailing during the Little Ice Age[31]. **E**. The probability of origin in West Africa based on enamel ^87^Sr/^86^Sr ratios and δ^18^O values, calculated and visualized for the donkey specimen from Jamestown. Highest probabilities of 80 to 90% can be found in coastal Senegambia and Sierra Leone. Presumably, the geology and climate of northern Senegal and Mauritania (not yet mapped for ^87^Sr/^86^Sr) would be an equally good fit.

### Trace metal concentrations

The maximum threshold concentrations (MTC) of key trace elements and rare earth elements were analyzed (Kamenov et al. 2018) for all samples assayed (Supplementary Material Section 4). Although Vanadium (V), Uranium (U), and Thorium (Th) in some of the samples are elevated above the MTC and indicate some degree of diagenetic alteration, the rest of the elements are within the possible in-vivo range. A highly intriguing observation in the trace element data indicates a possible anthropogenic in-vivo exposure to Ni for the animals. Both the donkey and horse tooth enamel show zones with extremely high Ni concentrations, way beyond values expected from natural diet and/or even diagenetic alteration. The horse enamel shows Ni as high as 1,684 ppm, and the donkey shows the highest values (Ni=6,847 ppm; Table S2).

## Discussion

### Consumption of domestic horses

Archaeozoological and taphonomic results provide strong corroboration of the association between the analyzed assemblage and the “Starving Time” winter of 1609, showing previously undocumented levels of intensive processing and consumption of domestic equids at Jamestown. Results indicate that at least two adult horses were eaten, butchered, and cooked or boiled, with most elements split open to extract even the most minute nutritional resources including dental pulp. These findings mirror historic texts describing the Starving Time consumption of horses as a last resort. John Smith in his *Generall Historie* (1624) describes the dire situation of the colony after his October 1609 departure, writing based on the observations of a witness to the Starving Time, “as for our Hogs, Hens, Goats, Sheepe, Horse, or what lived, our commanders, officers, and salvages daily consumed them”, noting that even the skin of the horses was eaten.

### Use of horses for transport

Osteological data indicate that either during the animals’ life at Jamestown or prior, at least some of the first domestic horses at the Jamestown colony were used for transport. Fractures to the enamel of a lower second premolar suggest bridling, while an asymmetric cross-sectional profile for a metapodial suggests repeated load and activity patterns linked with transport. These findings align well with the historical counts of seven horses arriving in 1609, and with recent investigation of equestrian artifacts from early James Fort-period contexts, including the First Well and Kitchen Cellar, in which seven sets of tack were also recovered. The two limb fractures identified are not necessarily linked to transport, but might relate to confinement of animals during trans-Atlantic transit, or during daily life at Jamestown.

### Presence of juvenile horse and donkey

Archaeozoological analyses reveal the presence of previously undocumented domestic equids during the earliest years at Jamestown. Although both male and female horses were mentioned in first person accounts of the Third Supply that arrived at Jamestown in late summer 1609, the presence of a juvenile horse specimen shows that imported horses had successfully begun to breed and reproduce at the colony before their slaughter.

Most incredibly, archaeofaunal materials demonstrate the previously unknown presence of a domestic donkey during the initial settlement of the Jamestown colony. Although donkeys were not mentioned in chronicles of the first voyages, John Smith included both horses and asses in a list of animals that would be likely to thrive in Virginia, based on the success of “them that were carried thither” suggests that Smith had, in fact, witnessed donkeys in the colony prior to his departure in 1609 [3]. Radiocarbon dates on this specimen are precisely consistent with measurements from other specimens in the feature, raising questions about where and how it might have reached this first settlement during the voyage.

### Heavy metal isotopes and concentrations

The Pb isotope ratios in the horse and donkey are distinct from what can be expected for North and South America, but fall within the observed range for historical European individuals (^206^Pb/^204^Pb=18.44 +/-0.10; ^207^Pb/^204^Pb=15.61 +/-0.04; ^208^Pb/^204^Pb=38.47 +/-0.19) exposed to anthropogenic Pb [21]. The presence of European anthropogenic Pb in the horse and donkey could either be due to “in-vivo” Pb exposure or post-mortem diagenetic changes. Although we cannot fully rule out in-vivo Pb exposure, the most likely explanation for the European anthropogenic Pb signature is contamination within the burial environment. Numerous European lead artifacts were recovered at the site. It is well-documented that when Pb artifacts are present in the burial environment their Pb isotopes quickly swamp any natural Pb signal present in buried teeth and bones [25]. Furthermore, a number of the enamel samples show elevated V, U, and Th concentrations, above the recommended Maximum Threshold Concentrations (MTC) for animal teeth [26], also providing evidence for post-mortem diagenetic alteration (Supplementary Material Section 4). Therefore, the most likely explanation for the observed “European Pb” signature is post-mortem contamination of the skeletal material with Pb released from the Pb artifacts.

The observed extremely high concentrations of Ni are unlikely to result from diagenetic alteration. Although no Ni concentrations and MTC are reported in Bertacchi et al. [26], in human enamel overall Ni is not highly enriched during taphonomic processes. For example, a compilation of diagenetically altered and unaltered archaeological human enamel shows maximum Ni values of 0.91ppm [27]. Similarly, analyses of modern faunal tooth enamel shows low Ni concentrations ranging between 0.1 and 2ppm [28]. In contrast, modern human enamel shows elevated Ni measurements, as high as 36.5ppm, due to the overall higher exposure to anthropogenic metals at present [27]. Although this is observed in modern human enamel, we can hypothesize that Ni will bioaccumulate in those animals exposed to high Ni sources. Potentially the most likely explanation for the observed extremely high Ni concentrations in the horse and donkey teeth is Ni exposure from the bits. Horse bits contained elevated Ni in the past, in some cases even bits were made of pure Ni, a practice less common at present due to the introduction of stainless steel bits after the 1940s [29]. Therefore, based on the observed high Ni in the enamel samples, most likely both animals were using high-Ni bits.

### Biomolecular evidence for trans-Atlantic exchange

Various lines of isotopic and genetic data allow us to exclude certain regions of origin for the two equids closely examined with biomolecular methods in this study. Isotope data may be partly contradictory, as isotopic characteristics overlap between geographic regions and most parts of the world are still not mapped for the complex isotopic variation found in the environment [30].

The analyzed horse sample has a δ^18^O value indicative of a colder climate than reported for present-day Britain [22]. This is particularly puzzling as the C_3_-fodder this equid received is what we would expect for the temperate climate of the United Kingdom. This kind of mismatch between the enamel δ^18^O values of archaeological horses and predictions for δ^18^O in modern precipitation were recently reported for horses from a medieval site in London [31], suggesting that the modern day meteoric δ^18^O values do not reflect the drinking water values of the historical past. In that study, Pryor and colleagues [31] associate this discrepancy with the Little Ice Age glaciation from ∼1550 to ∼1900 during which substantially colder climate affected the northern hemisphere.. If we adjust the δ^18^O values for an overall colder climate accordingly, the donkey clearly could not have come from Britain (see Figure 4), but the horse values may be consistent. Additionally, the horse’s ^87^Sr/^86^Sr and ^206^Pb/^204^Pb large match the isotope distribution of Britain), suggesting that it may indeed have been bred there and was later transported to the Americas via the Atlantic route.

The donkey’s δ^18^O values suggest provenience in warmer climates overall, fitting particularly well into δ^18^O isoscape predictions for the Mediterranean, including the Iberian Peninsula and North Africa [24] as well as the Caribbean [32]. This is supported by the mixed C_3_ and C_4_ dietary signal in the donkey specimen, suggesting that the fodder of this animal derived from a drier, warmer environment, home to wild and domesticated C_4_-grasses, including maize, sorghum, or millet, which were likely completely absent in 16^th^ and early 17^th^ century Britain. Hence, both the δ^18^O and δ^13^C data suggest possible origins for the donkey along the trans-Atlantic voyage route.

If we return to the consideration of the mobility of the Jamestown donkey specimen, its genetic ancestry provides important information hinting towards West Africa and the wider trade network associated with this region, all the way to the Iberian Peninsula. Strontium isotope ratios vary greatly in West Africa (0.709 to 0.730 [23]) and the Iberian Peninsula (0.7046 to 0.735 [33,34]), though less so in the Caribbean (0.706 to 0.711 [35]), but unfortunately no bioavailable ^87^Sr/^86^Sr data are yet available for North Africa. The donkey’s ^87^Sr/^86^Sr ratios of 0.7100 exceed the range of published Sr values from the Canaries, which tend to lower values from the islands’ young volcanic geology [36,37]. Similar values are rare in the Caribbean, outside of Trinidad, and are only found in biotic samples on metamorphic rocks in small parts of the islands, but more common in southern Portugal and to a lesser extent in western Spain. A novel strontium isoscape of West Africa allows us to model the probability for the donkey’s provenance to lie in West Africa at large, and we find high probabilities of origin along the coast of Senegambia and Sierra Leone (Figure S2). Although it has not yet been mapped for bioavailable ^87^Sr/^86^Sr, Mauritania, dominated by the same Cenozoic/Quaternary sediments as Senegal, is likely to have similar ^87^Sr/^86^Sr as Senegal, matching those of the Jamestown donkey. Given the known trajectory of the Third Supply voyage (Figure 1), plausible scenarios for the donkey’s origin include trade for an animal raised in mainland West Africa during a documented stopover in the Canaries, acquisition during an undocumented stop along the Iberian coast, or acquisition of a non-local animal from a more distant point of origin (e.g., Trinidad) during transit through the Caribbean.

When considered alongside genomic data showing a blend of west African and Iberian ancestry, these data appear to mirror patterns emerging from other taxa, such as cattle, showing a substantial contribution from African sources in the early colonial fauna from the Americas [38], and point towards a donkey drawn from emerging multicontinental and multicultural trans-Atlantic trade networks, blending populations from Africa and Europe into complex biological and economic networks.

### Ecological context of donkey demand and exchange at Jamestown

Historical correspondence and physical artifacts suggest that demand for donkeys at the colony, observed through the identification of unrecorded donkey in Starving Time contexts that likely emerged during trans-Atlantic travel, may have been anchored in deeper ecological processes. In comparison to horses, which are better adapted to colder and more arid grassland conditions found at higher latitudes, donkeys are an equatorial animal, with a likely ancestral home in Northeast Africa, and with their largest contemporary populations located in warmer climes [19,39]. In comparison to horses, donkeys consume significantly less forage and are more efficient with water, and in the contemporary United States, the highest donkey populations are found in the lower latitudes of the American South [40].

After the Starving Time, little to no documentary evidence records horses or donkeys being present in the colony. Despite this, martial law instituted in the spring of 1610 to restore order stipulated that the killing of a mare or horse was punishable by death [41]. In a June 1613 letter to England, Thomas Dale mentioned that the provisions of the colony should be sufficient, “provided always that we have beasts to manure the land, either horses or asses, but indeed the latter is best for us so they be great strong ones.” Later in the same letter he reminds the Virginia Company of his frequent requests for “100 she-asses” and several horses, which would be useful as work animals to plow the ground and pull carriages [42]. The emerging preference for donkeys over horses represents a departure from traditional English agricultural lifeways, which were best adapted for temperate landscapes and initially resulted in preferential success in places that recapitulated the climate of the British Isles [43,44].

Together, these patterns may reflect an emerging adaptation of domestic fauna to both the climate of colonial Virginia and the limitations of early foodways, in which conditions of habitat and food supply were favorable for donkey use but could not easily support horse rearing in early colonial contexts on the east coast, in contrast to the rapid speed with which horses spread into the continental interior of North America via the southern Great Plains and Rocky Mountains [2].

## Conclusion

Interdisciplinary scientific study of faunal remains associated with the earliest English settlement of the eastern seaboard of the United States shows that, consistent with historic records, domestic horses were transported across the Atlantic from the British Isles where they were used for transport and successfully reproduced before meeting their end in the Starving Time winter of 1609. Modification of these horse remains shows that extraordinarily intensive processing was used to survive this winter, including butchering, cooking, and extracting marrow and pulp from nearly all the equids found at the site. New discoveries of donkey remains demonstrate a new and previously unknown dispersal of domestic donkeys into the colony, likely involving undocumented trade with emerging trans-Atlantic networks of biological exchange linking Europe, Africa, and the Americas via the Caribbean. These results demonstrate the extraordinary diversity of emerging trans-Atlantic trade networks in livestock during the early 17^th^ century, and demonstrate the impact of animal ecology in shaping the early distribution of domestic livestock in the Americas.

## Materials and Methods

### Osteological study

To understand the role of domestic equids during the initial settlement of Jamestown, we conducted detailed osteological, taphonomic, and paleopathological study of 63 out of the 77 specimens morphologically confirmed as members of the *Equus* genus. For each specimen, we first identified the animal to taxon based on size and morphological comparisons to reference data from specimens in the Archaeozoology Laboratory at the University of Colorado-Boulder. For specimens with intact epiphyses or visible dentition, we assessed the likely age of each identified specimen using eruption and wear tables from Levine [45] and Evans et al. [46], and for cheek teeth, estimated age using crown height tables [45] and a Mitutoyo digital caliper. Although sex estimation is also sometimes possible based on pelvic morphology or the presence of intact mandibles and maxillae, the fragmented nature of the assemblage did not permit sex estimation in the assemblage.

### Taphonomy

For each specimen, we assessed life history, health and pathology information when possible, including evidence for fractures and disease [47], transport-linked damage to the dentition [12], and changes to the internal structure of the limbs [14,15]. We also assessed each specimen for evidence of nonhuman taphonomic processes such as weathering, root etching, and carnivore gnawing, as well as for cultural modifications including cut and chop marks, spiral fracturing, burning, pot polish, and other indicators of human activity. Each likely cut mark was also assessed under 20x and 200x microscopy using a DinoLITE digital microscope, and marks lacking a distinct “V” shaped profile were excluded from the analysis.

### Radiocarbon dating

We selected five equid tooth specimens for direct radiocarbon dating using Accelerator Mass Spectrometry (AMS) at the Keck-AMS facility at the University of California-Irvine, of which three produced successful measurements (Supplementary Material Section 2). Before conducting destructive analysis, each specimen was scanned using an EinScanSE structured light scanner to produce a 3D model of morphological data. We then prepared a simple Bayesian uniform phase model estimate for horse activity at the initial Jamestown colony following the methods of Ramsey [48], using an initial bounding date of the arrival of the Ponce de León expedition on the American mainland in 1513, assumed to be the first possible date for the arrival of domestic horses on mainland North America.

### Stable isotopes

A suite of isotopic analyses were conducted on two specimens from the Jamestown, Virginia site: a horse (JR2718K, #117576), and a donkey (JR3081F, #121161). After each tooth was thoroughly cleaned and excess plaque and residues mechanically removed, sequential samples of tooth enamel were drilled parallel to the growth axis, in the Bone Chemistry Laboratory in the Department of Anthropology, University of Florida.

Average data are provided below, and in the table (Supplementary Material Section 4). All data are from tooth enamel structural carbonate and capture that point in time during the mineralization of each specimen. Two sets of analyses were conducted in the Department of Geological Sciences, University of Florida: (1) Light isotope analysis on non-pretreated tooth enamel (structural carbonate) was measured by isotope ratio mass spectrometry (IRMS) with carbon (δ^13^C) and oxygen (δ^18^O) data; (2) Radiogenic isotope analysis on these same tooth enamel powders was measured using multiple collector-inductively coupled plasma mass spectrometry (MC-ICP-MS) for strontium (^87^Sr/^86^Sr) and lead (^206^Pb/^204^Pb) (Supplementary Material Section 4).

### DNA

A total of five equid archaeological remains were analyzed for DNA content at the ancient DNA facilities of the Centre for Anthropobiology and Genomics of Toulouse (CAGT, France). The experimental procedure followed the work from Librado et al. [49] and Todd et al. [19] (Supplementary Material Section 5). DNA screening revealed excellent DNA preservation in the donkey jack sample labeled JT02/121161 (labeled ‘Jamestown’ in Supplementary Appendix 1, and Figure 3, S3-S6). We then conducted phylogenetic analysis using Neighbor-Joining and TreeMix analyses, and explored the relationship of this specimen to global modern and ancient population structure using PCA, Struct-f4, and qpAdm (Supplementary Material Section 5). The sequence data generated in this study are available for download on the European Nucleotide Archive (Accession nb. XXX [to be added upon final submission]).

## Acknowledgments

This research was funded through an award by the National Science Foundation (NSF #1949305, “Horses and Human Societies in the American West”), and by France Génomique National infrastructure, funded as part of “Investissement d’avenir” program managed by Agence Nationale pour la Recherche (ANR-10-INBS-09); the CNRS International Research Project AnimalFarm; the France Génomique Appel à Grand Projet (ANR-10-INBS-09-08, MARENGO project), and; the European Research Council (ERC) under the European Union’s Horizon 2020 research and innovation programme (grant agreements 681605-PEGASUS, and 101071707-HorsePower). Special thanks to Jason Curtis (UF) for light isotope analysis and Dr. John Southon at UCI-Keck AMS facility for radiocarbon dating.

## Notes

### Competing Interest Statement

The authors have declared no competing interest.

