## Supplementary Material for "Early transatlantic movement of horses and donkeys at Jamestown"

**Section 1. Historical records from Jamestown.**

Few writings exist from the 1607-1619 period at Jamestown, and of those that do, the majority are first person accounts. These observations of the colony are often biased and incomplete, and thus must be considered carefully. The addition of archaeological finds and data to the information represented in the historic documents provides a clearer picture of life in early 17th century Virginia, but can also perpetuate questions and assumptions which may never be clarified. In this case, as represented in this manuscript, evidence strongly suggests that by the time the Starving Time winter was coming to a close in April 1610, all English-imported horses, mares, and donkeys in the colony had been killed for food. In addition to the writings of George Percy and John Smith, as mentioned in the manuscript, documentation of the colony written after the winter of 1609-1610 supports this.

Both Secretary William Strachey and Governor Lord De la Warr record a June 12, 1610 meeting called by the Governor during which he establishes a council and discusses pressing issues that need resolution, the first of which is obtaining and providing "provisions of victuals". As written by Strachey and published in 1612, he records the state of Jamestown just after the Starving Time winter: "It did not appear that any kind of flesh, deer, or what else of that kind could be recovered from the Indian, or to be sought in the country by the travail or search of his people; and the old dwellers in the fort, together with the Indians not to friend, who had the last winter destroyed and killed up all the hogs, insomuch as of five or six hundred (as it is supposed) there was not one left alive; nor an hen nor chick in the fort; and our horses and mares they had eaten with the first; and the provision which the lord general and captain general had brought, concerning any kind of flesh, was little or nothing, in respect it was not dreamt of by the adventurers in England that the swine were destroyed" [42].

There is no record of equids on any ships arriving at Jamestown after Gabriel Archer's 1609 mention of seven animals aboard the *Blessing* until at least 1614. Also notable is the distinct lack of horse or donkey bones recovered archaeologically from contexts which post-date the 1610 cleansing instituted by Lord De La Warr upon his arrival to the colony. For a human population estimated to be in the hundreds and continually growing, this relatively small number of horses during Jamestown's first years, combined with little artifactual evidence of horse riding in Jamestown's earliest archaeological contexts, suggests a cultural shift for the English colonists.

In England, ownership of a horse and the ability to travel was not only useful but indicative of social power. While a large assemblage of spurs at Jamestown indicates that the colonists clung to their English understanding of status signaling through fashion, the relative lack of horse furniture and tack, and small numbers of horses in the documentary and archaeological record, suggests that horses to be ridden were not a priority for the colonists in the first quarter of the seventeenth century. The colonists were adapting to a different life in Virginia, where large, strong animals were preferred for survival; for their usefulness to farming, and in times of desperation, as food [50].

Despite this, in May 1611, construction of "a stable for our horses" was recorded by Thomas Dale. Perhaps the colonists hoped that life in Virginia would begin to emulate what they were used to in England. Later that year, an Englishman taken prisoner by the Spanish at Point Comfort mentions "17 mares and horses" in the colony, although considering the circumstances, this may have been false information [42].

With this and future research in mind, it is perhaps interesting to consider later documentation for the import and breeding of mares, horses, and donkeys in Virginia. Nearly five years after the Starving Time winter in late April 1614, Don Diego de Molina, a Spaniard held prisoner at Jamestown wrote of horses captured by Captain Samuel Argall during a raid at Saint Sauveur, a French settlement on the coast of Maine. Molina records that the horses, along with other provisions and fifteen people from the French settlement were brought to Virginia. In that same year, Secretary of the colony Ralph Hamor records a thriving population in the colony of "some mares, horses, and colts", which perhaps included those captured by Argall, suggesting that the horse population, through import and/or local breeding and controlled by martial law was only beginning to increase[42].

As stated in the manuscript, Thomas Dale recognized the importance of horses and donkeys as work animals for the colony as early as 1613. However, it was not until 1616 that horses were again discussed in the same way. John Rolfe, writing A True Relation of the State of Virginia while in England with his wife, Matoaka (Pocahontas), indicates that “cattle, horses, mares, and goats” are carefully preserved for increase in the colony, and in a margin note, he specifies that the horses and oxen are a main want of the colonists because of their usefulness to pulling carts and plowing the ground for growing corn. Notably, Rolfe also records only 3 horses and 3 mares were present in the colony as of the spring of 1616, further evidence that horses played a different role in life in Virginia than they did in England in the first quarter of the 17th century [42].

**Section 2. Donkey identification.**

The occlusal surface of the upper M2 (catalog # 121161) identified as domestic donkey was sufficiently well preserved for further assessment. This tooth shows dental features that are typically found in *Equus asinus*. In particular, the distal length of the protocone is approximately equal to the mesial length of the protocone (i.e., the protocone is symmetrical), the pli caballin is absent, and the post-protoconal groove is deep. These features are found at a higher frequency in domestic donkeys than in domestic horses in upper M1 and M2 in middle stages of wear (i.e., excluding teeth in early and late wear; CIB-O unpublished data): *E. asinus* with symmetrical protocone (M1 = 92.3%, M2 = 75.0%, n = 13, 12), pli caballin absent (M1 = 92.3%, M2 = 75.0%, n = 13, 12), deep post-protoconal groove (M1 = 46.2%, M2 = 75.0%, n = 13, 12); *E. caballus* with symmetrical protocone (M1 = 0.0%, M2 = 11.1%, n = 18), pli caballin absent (M1 = 44.4%, M2 = 33.3%, n = 18), deep post-protoconal groove (M1 = 5.6%, M2 = 61.1%, n = 18). The presence of all three characters (symmetrical protocone, pli caballin absent, and deep post-protoconal groove) in the same tooth is also found at a higher frequency in domestic donkeys than in domestic horses (*E. asinus*: M1 = 46.2%, M2 = 50.0%, n = 13, 12; *E. caballus*: M1 = 0.0%, M2 = 0.0%, n = 18).

**Section 3. Radiocarbon dating.**

Preparation of bone and tooth samples followed Shammas et al. [51]: briefly, following mechanical surface cleaning if required, samples of 200 mg of bone or dentin were crushed to ~1 mm powder and decalcified overnight in a measured amount of 1N HCl calculated as just sufficient to dissolve the entire sample if all of the material was hydroxyapatite. They were then washed with MQ water, gelatinized overnight at 60°C in 0.01N HCl, ultrafiltered in precleaned Vivaspin 15 devices to select a high molecular weight fraction (>30kDa), and freeze dried.

Samples of 1.5 - 2 mg of pretreated organics were combusted with CuO and Ag wire getter at 900°C in quartz tubes sealed under vacuum, graphitized by hydrogen reduction at 525°C with an iron powder catalyst (Santos et al., 2007), and the Fe-graphite mixture was pressed into Al sample holders. Radiocarbon was measured at the Keck Laboratory at University of California Irvine on a National Electrostatics 1.5SDH Compact AMS system [53]. Radiocarbon ages are shown as conventional $^{14}C$ ages corrected for isotopic fractionation, with 1 sigma uncertainties that reflect scatter in repeated runs and uncertainties in the measurement of blanks and normalizing standards (NIST Oxalic Acid 1, SRM4990B) as well as counting statistics. Aliquots of ~0.7mg of ultrafiltered collagen for stable isotope ratio and elemental (%C and %N) analyses were placed in tin capsules and measured using a Fisons NA 1500NC elemental analyzer and Finnigan Delta Plus isotope ratio mass spectrometer, at precisions of ±0.1‰ and ± 0.2‰ for $\delta^{13}C$ and $\delta^{15}N$, respectively.

| **Lab # (UCIAMS)** | **Catalog #** | **$\delta^{13}C$ (‰)** | **±** | **fraction** | **±** | **$\delta^{14}C$ (‰)** | **±** | **$^{14}C$ age (BP)** | **±** |
|---|---|---|---|---|---|---|---|---|---|
| 271743 | 121161 | -19.2 | 0.1 | 0.9537 | 0.0018 | -46.3 | 1.8 | 380 | 20 |
| 271744 | 121460 | -22.7 | 0.1 | 0.9545 | 0.0018 | -45.5 | 1.8 | 375 | 15 |
| 271745 | 117294 | -23.2 | 0.1 | 0.9558 | 0.0017 | -44.2 | 1.7 | 365 | 15 |

*Table S1. Radiocarbon dates and associated quality control data from three successfully dated horse and donkey specimens.*

**Section 4. Isotope analysis.**

Sample processing for Sr and Pb isotope analysis was done in a class 1000 Clean Lab, equipped with class 10 laminar flow hoods, in the Department of Geological Sciences, University of Florida. Tooth enamel samples were dissolved in pre-cleaned Teflon vials in 8N $HNO_3$ (optima). The vials were then opened and evaporated to dryness in a laminar flow hood. Sr and Pb were separated by ion chromatography from single aliquots. The dissolved and dried residues were first dissolved in 1N Seastar HBr and passed through Dowex 1X-8 (100-200 mesh) resin to separate Pb for isotope analyses. During the lead elution step all of the wash cuts were collected for subsequent Sr separation, as the latter is not absorbed on the Dowex resin. The wash was dried down on the hot plate and the dried residues were dissolved in 3.5N $HNO_3$ and loaded on to cation exchange columns packed with Sr-spec resin (Eichrom Technologies, Inc.) to separate Sr for isotope analyses. Sr and Pb isotopic ratios were measured using a "Nu-Plasma" MC-ICP-MS. The results are reported relative to NBS 987 $^{87}Sr/^{86}Sr$=0.710246 (+/- 0.000030). Pb isotopic analyses were conducted using Tl normalization and the results are reported relative to NBS 981 $^{206}Pb/^{204}Pb$=16.937 (+/-0.004), $^{207}Pb/^{204}Pb$=15.490 (+/-0.003), and $^{208}Pb/^{204}Pb$=36.695 (+/-0.009).

To compare the equine $\delta^{18}O$ data to drinking and/or precipitation water in various parts of the world, we followed the same procedure and equations outlined for archaeological horses by Pryor et al. [31]. We first converted $\delta^{18}O$ carbonate measurements from the international standard VPDB to SMOW [54], then carbonate $\delta^{18}O$ values to phosphate $\delta^{18}O$ values [55], then phosphate $\delta^{18}O$ values to drinking water values, using two different equations developed for horses [56,57].

To map the equine $^{87}Sr/^{86}Sr$, $^{206}Pb/^{204}Pb$ and drinking water $\delta^{18}O$ data to Britain we relied on the interactive biosphere isotope database [22] and exported maps with permission. To map the probability of origin for the donkey specimen in West Africa using its $^{87}Sr/^{86}Sr$ ratios and drinking water $\delta^{18}O$ values, we employed a strontium isoscape of sub-Saharan Africa [23] and global oxygen isotope precipitation data [24] available at www.waterisotopes.org, together with the continuous-surface assignment framework from the R package "assignR" [58]. This analysis was performed in the R environment, version 4.3.1 (https://www.r-project.org/).

| taxon | specimen | tooth sampled | $^{87}Sr/^{86}Sr$ | $^{206}Pb/^{204}Pb$ | $\delta^{13}C$ VPDB | $\delta^{18}O_{carbonate}$ VPDB | $\delta^{18}O_{carbonate}$ SMOW | $\delta^{18}O_{phosphate}$^ SMOW | $\delta^{18}O_{water}$* SMOW | $\delta^{18}O_{water}$** SMOW |
|---|---|---|---|---|---|---|---|---|---|---|
| horse | JR2718K, #117576 | M3 | 0.7104 | 18.469 | -13.2 | -6 | 24.2 | 15.2 | -11.2 | -10.5 |
| donkey | JR3081F, #121161 | M2 | 0.7100 | 18.449 | -8.8 | -2.5 | 27.6 | 18.8 | -5.4 | -5.4 |

^ after Iacumin et al. [55]
* after Pederzani et al. [57]
** after Delgado Huertas et al. [56]

***Table S2.*** *Summary of average strontium, lead, carbon and oxygen isotope data from a Jamestown horse and donkey dental enamel. Enamel carbonate $\delta^{18}O$ values were converted to $\delta^{18}O$ precipitation/drinking water values using equations outlined by Pryor et al. [31]. Light isotope data are reported in ‰.*

***Table S3.*** *Trace element concentrations in ppm for analyzed samples.*

| | Horse (#117576) | | | | Donkey (#121161) | | | | |
|---|---|---|---|---|---|---|---|---|---|
| **Concentrations in ppm** | **117576.1** | **117576.3** | **117576.5** | **117576.7** | **121161.1** | **121161.3** | **121161.5** | **121161.7c** | **121161.7s** |
| Mg | 2188 | 2102 | 1964 | 2423 | 1981 | 2193 | 2197 | 2417 | 1909 |
| Al | 30.8 | 26.7 | 29.4 | 310.5 | 54.9 | 36.5 | 282.8 | 8.3 | 75.8 |
| Ca | 329639 | 318242 | 309075 | 321342 | 305768 | 319035 | 334264 | 333680 | 266888 |
| V | 2.611 | 2.164 | 4.144 | 1.704 | 4.186 | 2.493 | 3.375 | 1.472 | 1.837 |
| Cr | 1.688 | 1.775 | 2.374 | 8.101 | 11.353 | 18.453 | 4.778 | 0.113 | 1.140 |
| Mn | 47.332 | 27.714 | 20.513 | 80.303 | 52.188 | 47.964 | 132.952 | 33.431 | 45.889 |
| Fe | 76.250 | 63.564 | 73.100 | 453.992 | 270.562 | 294.832 | 610.537 | 133.134 | 188.554 |
| Ni | 1365.375 | 39.578 | 84.065 | 1683.856 | 164.264 | 6847.350 | 2161.163 | 0.157 | 511.792 |
| Cu | 3.197 | 2.963 | 3.649 | 7.740 | 1.682 | 2.934 | 3.484 | 3.042 | 2.442 |
| Zn | 46.634 | 44.807 | 52.025 | 45.149 | 48.774 | 51.865 | 95.568 | 43.092 | 70.770 |
| Rb | 0.645 | 0.319 | 0.401 | 0.979 | 0.501 | 0.500 | 0.868 | 0.443 | 0.718 |
| Sr | 457.411 | 505.666 | 644.227 | 548.904 | 644.090 | 647.616 | 540.295 | 347.188 | 349.773 |
| Ba | 59.570 | 53.580 | 71.941 | 67.905 | 88.605 | 46.175 | 78.096 | 36.378 | 37.585 |
| La | 0.188 | 0.200 | 0.238 | 1.019 | 0.436 | 0.597 | 1.418 | 0.408 | 0.834 |
| Ce | 0.226 | 0.158 | 0.224 | 1.108 | 0.320 | 0.429 | 1.465 | 0.188 | 0.362 |
| Pr | 0.152 | 0.083 | 0.107 | 0.276 | 0.161 | 0.202 | 0.357 | 0.149 | 0.245 |

| | | | | | | | | | |
|---|---|---|---|---|---|---|---|---|---|
| Nd | 0.092 | 0.103 | 0.148 | 0.665 | 0.314 | 0.468 | 1.094 | 0.257 | 0.545 |
| Sm | 0.029 | 0.023 | 0.036 | 0.141 | 0.070 | 0.099 | 0.221 | 0.062 | 0.098 |
| Eu | 0.061 | 0.031 | 0.041 | 0.073 | 0.057 | 0.063 | 0.086 | 0.052 | 0.081 |
| Gd | 0.050 | 0.039 | 0.059 | 0.154 | 0.103 | 0.131 | 0.241 | 0.084 | 0.155 |
| Tb | 0.162 | 0.076 | 0.096 | 0.144 | 0.121 | 0.136 | 0.140 | 0.121 | 0.184 |
| Dy | 0.051 | 0.039 | 0.053 | 0.129 | 0.086 | 0.117 | 0.201 | 0.071 | 0.130 |
| Ho | 0.148 | 0.070 | 0.089 | 0.134 | 0.112 | 0.126 | 0.130 | 0.112 | 0.172 |
| Er | 0.031 | 0.021 | 0.031 | 0.069 | 0.049 | 0.061 | 0.099 | 0.039 | 0.076 |
| Tm | 0.148 | 0.069 | 0.087 | 0.125 | 0.106 | 0.118 | 0.111 | 0.107 | 0.161 |
| Yb | 0.051 | 0.030 | 0.040 | 0.075 | 0.055 | 0.066 | 0.093 | 0.049 | 0.089 |
| Lu | 0.136 | 0.063 | 0.079 | 0.113 | 0.097 | 0.107 | 0.099 | 0.098 | 0.147 |
| Pb | 1.108 | 0.946 | 0.911 | 8.325 | 1.914 | 2.200 | 5.057 | 1.746 | 4.371 |
| Th | 0.161 | 0.077 | 0.096 | 0.194 | 0.119 | 0.127 | 0.256 | 0.112 | 0.173 |
| U | 0.423 | 0.255 | 0.301 | 0.193 | 0.221 | 0.233 | 0.614 | 0.286 | 0.736 |

| | MTC1 (ppm) | sample | 6197 | 6196.1 | 6196.2 | 6196.3 | 6146.2 11.1 | 6146.2 11.3 | 6146.2 11.5 | 6146.2 11.7 | 6147.11 1.1 | 6147.11 1.3 | 6147.11 1.5 | 6147.11 1.7c | 6147.11 1.7s |
|---|---|---|---|---|---|---|---|---|---|---|---|---|---|---|---|
| **V** | **1.8** | **V/MTC1** | 0.01 | 0.37 | 0.27 | 0.07 | 1.45 | 1.20 | 2.30 | 0.95 | 2.33 | 1.38 | 1.87 | 0.82 | 1.02 |
| **Fe** | **2510** | **Fe/MTC1** | 0.00 | 0.01 | 0.00 | 0.00 | 0.03 | 0.03 | 0.03 | 0.18 | 0.11 | 0.12 | 0.24 | 0.05 | 0.08 |
| **La** | **1.9** | **La/MTC1** | 0.02 | 0.13 | 0.08 | 0.05 | 0.10 | 0.11 | 0.13 | 0.54 | 0.23 | 0.31 | 0.75 | 0.21 | 0.44 |
| **Ce** | **7.8** | **Ce/MTC1** | 0.01 | 0.04 | 0.03 | 0.02 | 0.03 | 0.02 | 0.03 | 0.14 | 0.04 | 0.06 | 0.19 | 0.02 | 0.05 |
| **Nd** | **1.2** | **Nd/MTC1** | 0.01 | 0.20 | 0.13 | 0.07 | 0.08 | 0.09 | 0.12 | 0.55 | 0.26 | 0.39 | 0.91 | 0.21 | 0.45 |
| **Dy** | **0.4** | **Dy/MTC1** | 0.05 | 0.13 | 0.11 | 0.08 | 0.13 | 0.10 | 0.13 | 0.32 | 0.22 | 0.29 | 0.50 | 0.18 | 0.33 |
| **Yb** | **0.1** | **Yb/MTC1** | 0.21 | 0.36 | 0.30 | 0.25 | 0.51 | 0.30 | 0.40 | 0.75 | 0.55 | 0.66 | 0.93 | 0.49 | 0.89 |
| **Th** | **0.1** | **Th/MTC1** | 0.70 | 0.80 | 0.67 | 0.74 | 1.61 | 0.77 | 0.96 | 1.94 | 1.19 | 1.27 | 2.56 | 1.12 | 1.73 |
| **U** | **0.1** | **U/MTC1** | 0.42 | 0.85 | 0.59 | 0.55 | 4.23 | 2.55 | 3.01 | 1.93 | 2.21 | 2.33 | 6.14 | 2.86 | 7.36 |

***Table S4.*** *Trace element concentrations of the donkey and horse enamel normalized by the MTC1 (Maximum Threshold Concentrations) for all mammals. MTC1 from Betacchi et al. 2024. Element/MTC values above 1 indicate some degree of diagenetic alteration, note in some samples V/MTC1, Th/MTC1, and U/MTC1 are above 1.*

| UF BCL No. | δ13C (‰) | δ18O (‰) | O/C Lane | Distance from crown (mm) | Sr/Pb Lane | $^{87}Sr/^{86}Sr$ | ± | $^{208}Pb/^{204}Pb$ | ± | $^{207}Pb/^{204}Pb$ | ± | $^{206}Pb/^{204}Pb$ | ± | $^{208}Pb$ (V) |
|---|---|---|---|---|---|---|---|---|---|---|---|---|---|---|
| Horse (#117576) | | | | | | | | | | | | | | |
| 6146_211.01 | -12.7 | -5.6 | 1 | 0 | 1+2 | 0.710219 | (±0.000026) | 38.3955 | (±0.012) | 15.6364 | (±0.005) | 18.4455 | (±0.0057) | 0.5 |
| 6146_211.02 | -13.2 | -6.3 | 2 | 5 | | | | | | | | | | |
| 6146_211.03 | -13.4 | -6.3 | 3 | 10 | 3+4 | 0.710290 | (±0.000025) | 38.4282 | (±0.0032) | 15.6271 | (±0.0012) | 18.4861 | (±0.0013) | 3.1 |
| 6146_211.04 | -13.4 | -6.2 | 4 | 15 | | | | | | | | | | |
| 6146_211.05 | -12.6 | -5.6 | 5 | 20 | 5+6 | 0.710624 | (±0.000016) | 38.4487 | (±0.003) | 15.6295 | (±0.001) | 18.5036 | (±0.0012) | 6.1 |
| 6146_211.06 | -13.3 | -5.7 | 6 | 25 | | | | | | | | | | |
| 6146_211.07 | -13.5 | -5.8 | 7 | 30 | 7+8 | 0.710470 | (±0.00002) | 38.4043 | (±0.0019) | 15.6250 | (±0.00063) | 18.4438 | (±0.00068) | 5.8 |
| 6146_211.08 | -13.3 | -6.3 | 8 | 35 | | | | | | | | | | |
| Average | -13.2 | -6.0 | | | | 0.71040 | | 38.419 | | 15.630 | | 18.470 | | |
| Std. Dev. | 0.328 | 0.339 | | | | 0.0002 | | 0.024 | | 0.005 | | 0.030 | | |
| Donkey (#121161) | | | | | | | | | | | | | | |
| 6147_111.01 | -8.8 | -2.4 | 1 | 0 | 1+2 | 0.710141 | (±0.000022) | 38.4139 | (±0.003) | 15.6266 | (±0.0011) | 18.4615 | (±0.0013) | 4.8 |
| 6147_111.02 | -8.4 | -2.8 | 2 | 5 | | | | | | | | | | |
| 6147_111.03 | -8.3 | -2.6 | 3 | 10 | 3+4 | 0.709780 | (±0.000017) | 38.3666 | (±0.0022) | 15.6254 | (±0.00083) | 18.4101 | (±0.00094) | 5.6 |
| 6147_111.04 | -8.5 | -2.5 | 4 | 15 | | | | | | | | | | |
| 6147_111.05 | -9.2 | -2.2 | 5 | 20 | 5+6 | 0.710222 | (±0.000016) | 38.4344 | (±0.0024) | 15.6276 | (±0.0009) | 18.4643 | (±0.00097) | 4.2 |
| 6147_111.06 | -9.4 | -2.4 | 6 | 25 | | | | | | | | | | |
| 6147_111.07 | -8.9 | -2.8 | 7 | 30 | 7 | 0.710020 | (±0.000026) | 38.4322 | (±0.0024) | 15.6314 | (±0.00081) | 18.4609 | (±0.00094) | 3.6 |
| Average | -8.8 | -2.5 | | | | 0.71004 | | 38.412 | | 15.628 | | 18.449 | | |
| Std. Dev. | 0.434 | 0.229 | | | | 0.0002 | | 0.031 | | 0.003 | | 0.026 | | |

***Table S5.*** *Sequential isotope measurements from horse and donkey remains analyzed in the study.*

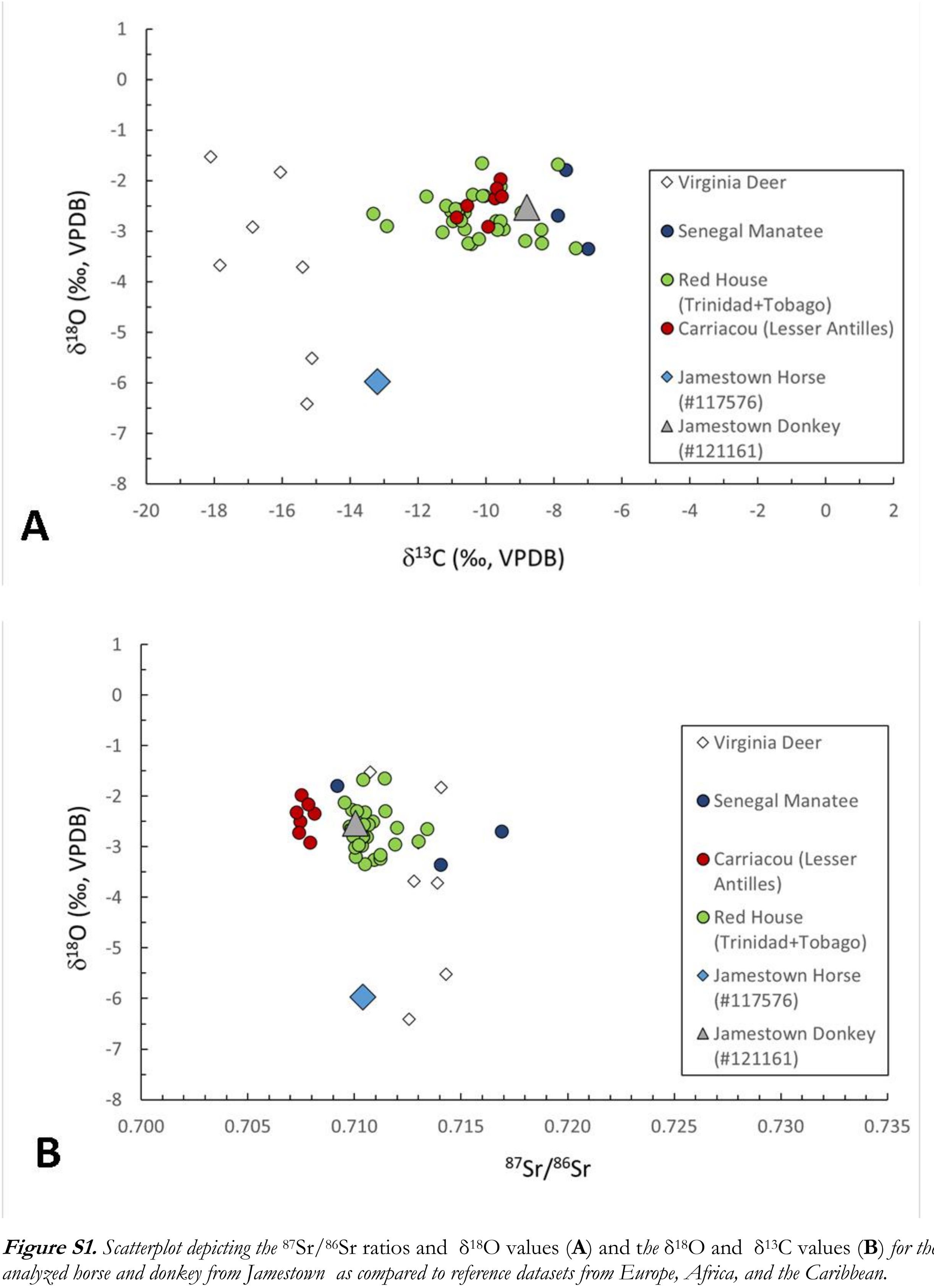


***Figure S1.*** *Scatterplot depicting the* $^{87}Sr/^{86}Sr$ ratios and $\delta^{18}O$ values (**A**) and t*he* $\delta^{18}O$ and $\delta^{13}C$ values (**B**) *for the analyzed horse and donkey from Jamestown as compared to reference datasets from Europe, Africa, and the Caribbean.*

**Section 5. DNA analysis.**

DNA extraction procedures, following the work of Librado et al. [49] and Todd et al. [19]involved: (1) bleaching 220-440 mg of osseous powder prior to DNA extraction, (2) powder pre-digestion for 45 mins at 37°C, (3) treatment of DNA extracts with the USER enzymatic mix, and; (4) preparation of triple-indexed DNA libraries compatible with Illumina paired-end DNA sequencing. DNA preservation levels were assessed from 357,723-737,059 MiniSeq sequencing reads (Table S1), following demultiplexing and read-collapsing with AdapterRemoval (v2.3.1, --barcode-mm-r1 1 --barcode-mm-r2 1 --minlength 25 --trimns --trimqualities --minadapteroverlap 3 --mm 5[59]), Bowtie2 mapping with local realignment against the donkey reference nuclear and mitochondrial (Genbank Accession nb. NC_016061) reference genomes, and filtering of low-quality and repeated alignments (mapping quality Phred-scores < 25). Those post-sequencing read processing steps were carried out using the automated bioinformatic pipeline Paleomix (v1.2.13[60]), with the parameters recommended by Poullet and Orlando [61]. High-quality unique read alignments against the horse EquCab2 nuclear [62]and mitochondrial (Genbank Accession nb. NC_001640) were obtained following the same procedure, and were used as input for sex determination and species identification with Zonkey (v1.2.13 [63] Table S1).

A total of 4 DNA libraries were prepared for deeper sequencing on the Illumina NovaSeq6000 instrument (S4 lane), which provided a total of 205.2 million sequencing reads, including 188.4 million pairs that were collapsed using AdapterRemoval2 on the basis of the significant sequence overlap identified between read mates (Table S1). These data provided a characterization of the genome sequence of the JT02 specimen at an average 0.54-fold depth-of-coverage (Table S1). The BAM alignment profile was combined to a comparative genome panel including 240 additional donkeys, of which 32 ancient donkeys [19], and seven Tibetan kiangs (*Equus kiang*), four wild onagers (*Equus hemionus*), and three zebras (*Equus burchelli boehmi*, *Equus grevyi*, and *Equus zebra hartmannae*), used as outgroups. These genomes were characterized following the same read processing and mapping procedures as for specimen JT02, downloading sequencing data from public repositories [64]. As the genome coverage achieved for the JT02 specimen (0.54-fold) was inferior to the ~0.75-fold minimal requirements characterized by Todd and colleagues [19] for imputing genotypes, the data were pseudo-haploidized, following the methodology from Taylor et al. [2], tolerating 80% missingness at most, and requiring base quality Phred scores of at least 30. This resulted in the production of a tped matrix of single nucleotide polymorphisms, containing 10,774,885 autosomal sites in 255 individuals.

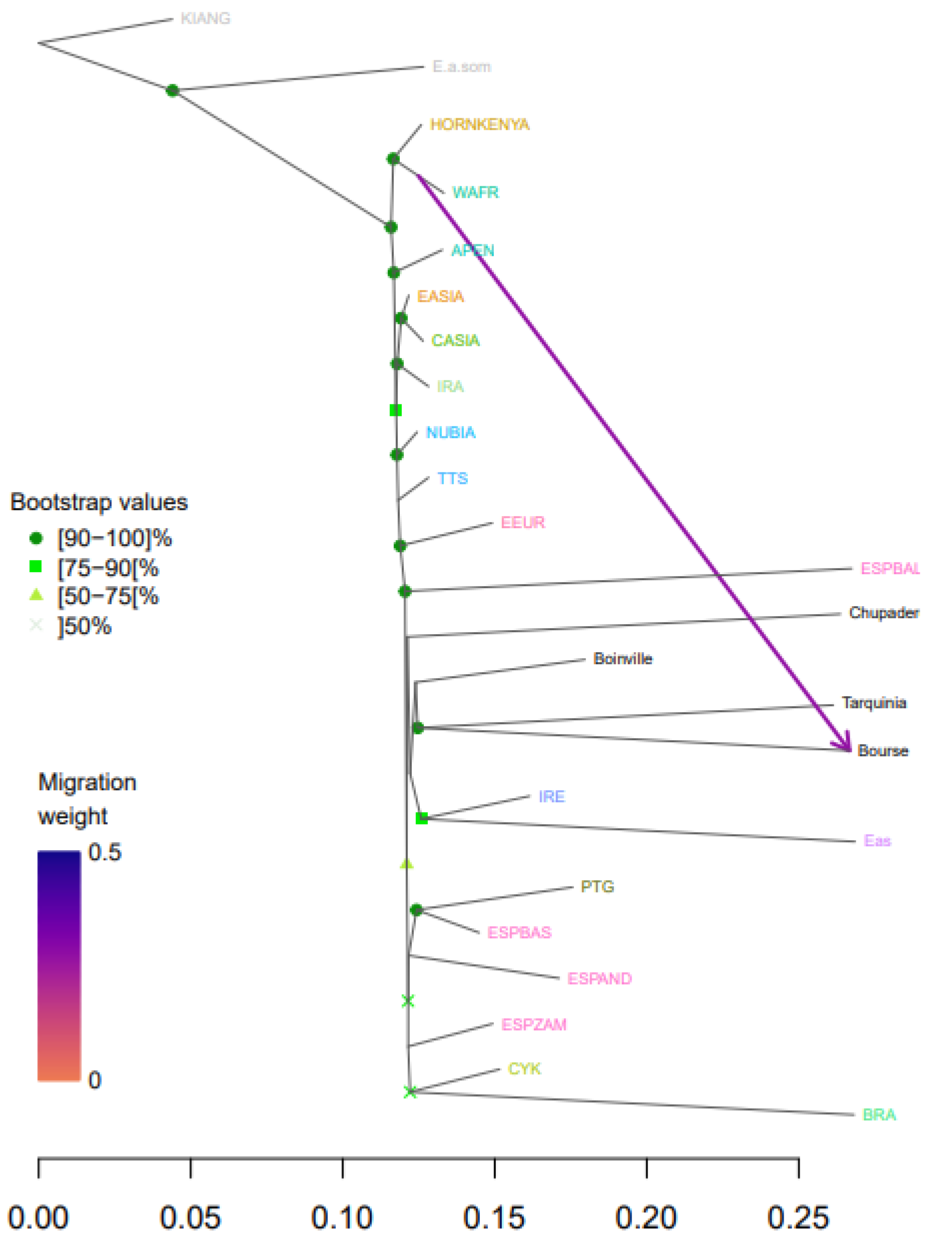


***Figure S2***. *Phylogenetic relationships between the ancient donkey specimen CHUPADEROS, and three ancient groups of donkeys from France (Boinville, 200-500 CE, and; Bourse, 0-500 CE) and Italy (Tarquinia, 803-412 BCE), as well as a worldwide panel of modern donkeys, stratified by subcontinental groups of genetic affinities. The tree was rooted on the sequence data available for the Equus kiang species, and node supports were assessed from 100 bootstrap pseudo-replicates. The tree models population genetic affinities, considering an optimal number of migration edges (Table S2). Subcontinental groups of genetic affinities are defined according to Todd et al. [19], with Spanish accession subdivided to gain further resolution into possible genetic affinities in the region (ESPAND: Andalusia; ESPBAS: Basque; ESPZAM: Zamorano Leones, and; ESPBAL: Baleares).*

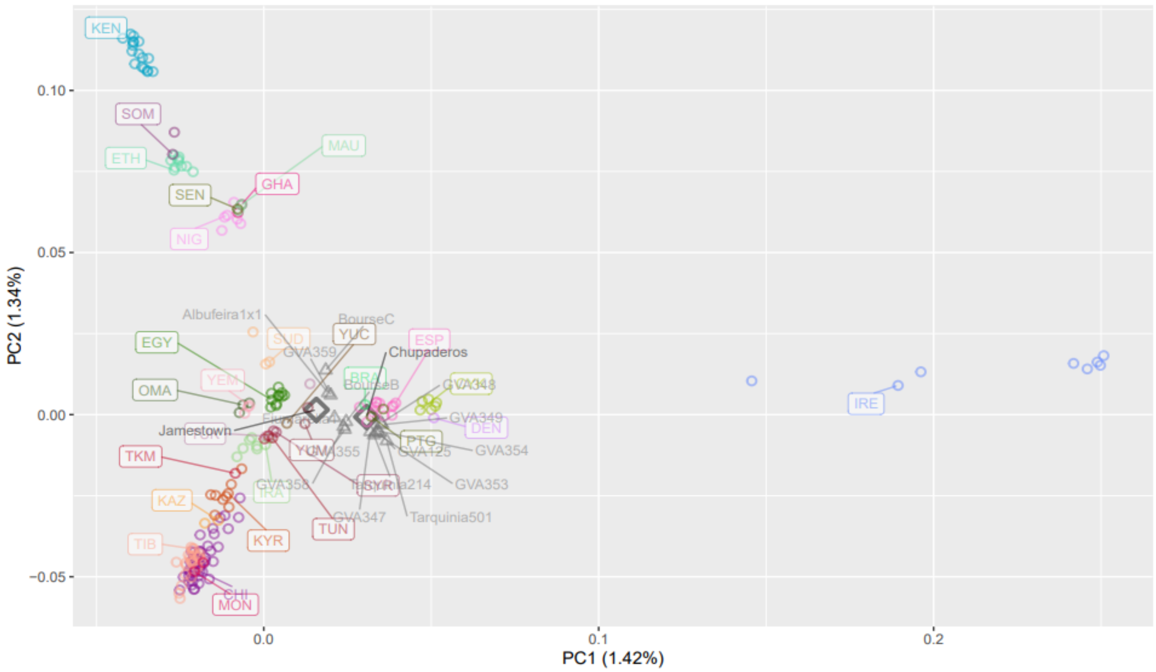


***Figure S3.*** *Principal Component Analysis revealing worldwide genetic affinities of donkey populations. A) Principal components 1 and 2. For clarity, we label only one of the modern accessions from a given country. The color correspondence is provided in Supplementary Fig. S5.*

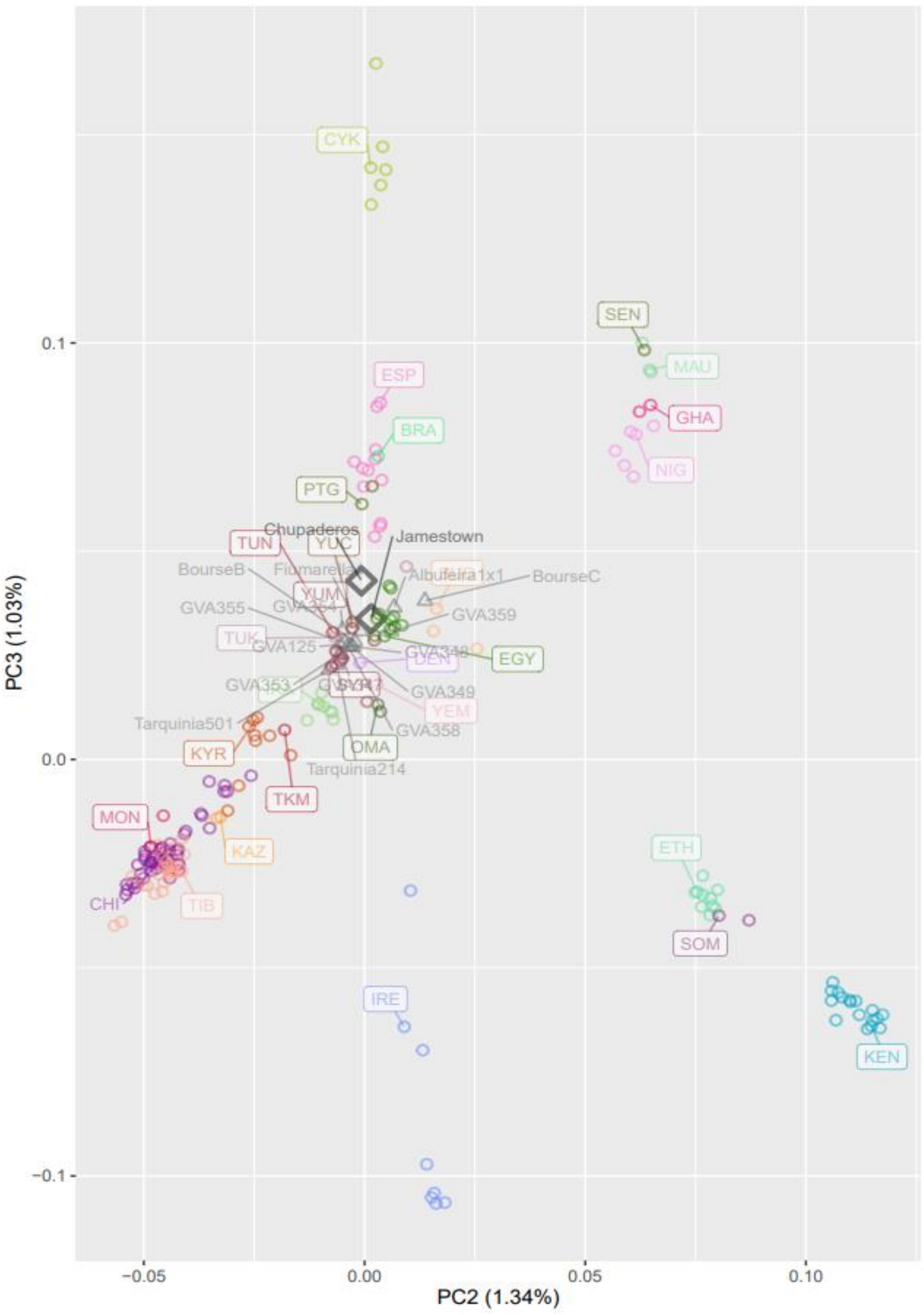


***Figure S4.*** *Principal components 2 and 3. The percentage reported on each axis indicates the fraction of the overall variance explained by the corresponding component. For clarity, we label only one of the modern accessions from a given country. The color correspondence is provided in Supplementary Fig. S5.*

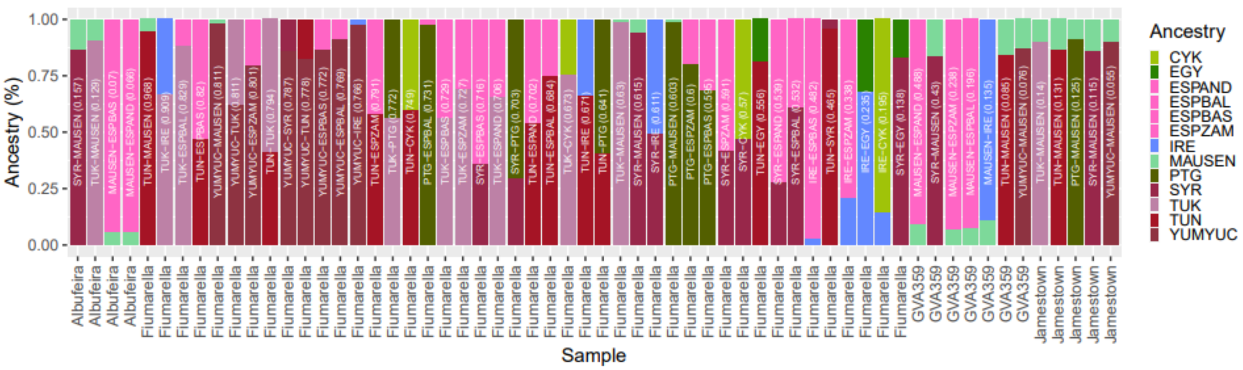


***Figure S5.*** *qpAdm 2-way modelling of the genomic makeup of specific ancient donkey specimens. The ancient individuals or groups of individuals (GVA2) are the same as those considered in Fig. 6. The barplots indicate the respective admixture contributions of the two population sources for which 2-way qpAdm modelling was not rejected (the corresponding p-value associated to each model is provided between parentheses).*

The genetic variation present in the matrix was processed to reconstruct Neighbour-Joining (NJ) phylogenetic affinities, assessing node robustness from 100 bootstrap pseudo-replicates. First, a NJ tree was obtained applying FastMe (v2.1.6.2[65]) to the pairwise genetic distances calculated using Plink v1.9 (v1.90b6.24; --distance square 1-ibs flat-missing) on the complete tped matrix. This analysis (not shown) served to confirm the placement of the outgroups. Then, the same analysis was rerun excluding the 14 outgroups, and 18 modern donkeys demonstrated by Todd and colleagues [19] to be associated with a complex genomic makeup and/or to have been translocated outside their native regions. The resulting tree was rooted manually (Figure 3), according to the placement of the outgroups indicated in the first analysis. Phylogenetic affinities were further explored using TreeMix (v1.13[66]) and unlinked SNPs only, pruning sites in linkage disequilibrium while requiring minimal allelic frequencies of 5% in Plink v1.9 (--maf 0.05 –indep-pairwise 500 10 0.2) and keeping *Equus kiang* specimens as outgroups. These analyses were based on 180,007-554,460 sites (Table S2). They grouped the modern donkey genomes into the population groups defined by Todd and colleagues [19], and added ancient individuals, or groups of ancient individuals forming subgroups close-kin relatives (at Boinville-en-Woëvre, GVA1 And GVA2; Table S2), one at a time to assess their individual placement (grouping modern accessions by country of origins, and grouping modern accessions according to the subcontinental groups of genetic affinities defined by Todd and colleagues [19]. The optimal number of migration edges was determined following Todd et al. [19] using the mixed linear model implemented in the optM R package. Node support was assessed following the procedures described as part of the BITE package [67].

A principal component analysis (PCA) was carried out to assess the worldwide donkey population structure and those groups genetically closest to the JT02/121161 specimen, using the matrix of pruned DNA variants and considering 5% as a minimal allelic frequency (Figure S3,S4). This analysis was carried out using smartPCA [68] for projecting ancient specimens onto the population structure present in modern populations (inbreed: YES, autoshrink: YES, lsqproject: YES). Population genetic affinities were also assessed from f3-Outgroup and f4-statistics (Fig. 3[69]), which were calculated using the Calc-f3 and Calc-f4 functions distributed as part of the Struct-f4 statistical package from Librado and Orlando [70]. The procedure was applied to the matrix of linked genetic variants, using *Equus kiang* as outgroup and grouping modern accessions by countries of origins, or the subcontinental groups of genetic affinities defined by Todd and colleagues [19]. Finally, the qpAdm package from Admixtools (v6.5.1) was used to model the genomic makeup of the same specimens as members of predefined groups of modern donkey populations, considering models of 1-way and 2-ways admixture (allsnps: NO, inbreed: NO). Based on the population affinities revealed in the analyses presented above, qpAdm analyses considered *Equus kiang*, SOM, KEN, ETH, SOM, IRA, KYR, KAZ, CHI, TIB, MON, YEM, OMA, SUD, NIG and GHA populations as outgroup *'Right'* populations. All other combinations of 'left' populations, including one ancient specimen and the remaining modern accessions (grouped by country of origins) were considered. Those associated with p-values >5% were retained as plausible models for the genomic makeup of the ancient individual considered (Figure S5).

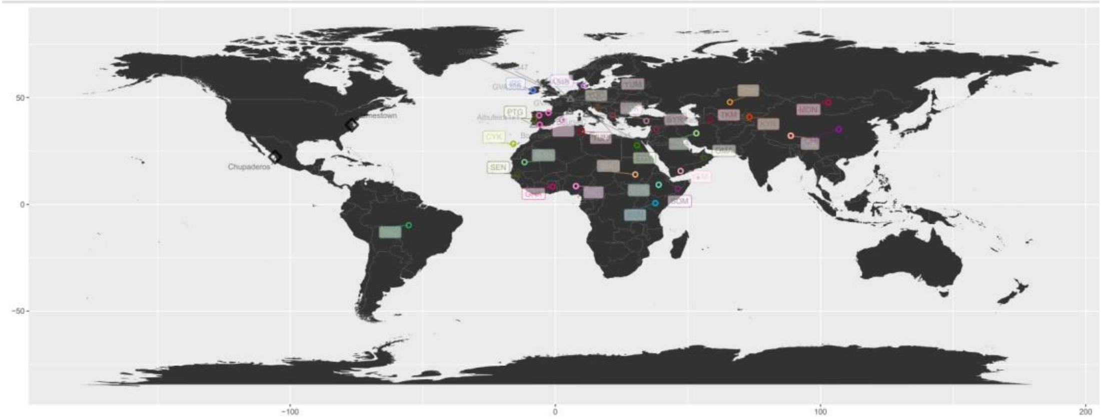


***Figure S6.*** *Geographic distribution of the 241 donkey genomes analyzed in this study. The comparative panel was originally published by Todd and colleagues (Todd et al. [19], and consisted of 208 genomes from modern donkeys spread across the world. It also included a complement set of ancient donkey genomes, of which 31 from Europe were included in this study (colored in Grey). The donkey genome data released by Taylor and colleagues (Taylor et al. [2], characterizing the genome of an ancient donkey specimen from Chupaderos, Mexico, at an average 1.65-fold average depth-of-coverage, were also included.*

***Supplementary Appendix***. *DNA samples analyzed in this study and associated metadata.*
